# Characterizing *Plasmodium falciparum* genetic diversity and complexity of infections in clinical malaria infections in Western and Coastal Kenya using the poly-alpha microsatellite marker

**DOI:** 10.1101/2025.04.26.650801

**Authors:** Nicole Achieng’, Kelvin M. Kimenyi, Cavin Mgawe, Leonard Ndwiga, L. Isabella Ochola-Oyier

## Abstract

**Background:** Genotyping *P. falciparum* polymorphic merozoite genes to describe parasite genetic diversity and the complexity of malaria infections (COI) is routinely used to assess the effectiveness of malaria control interventions. They are also utilized in anti-malarial drug therapeutic efficacy studies (TES) to differentiate recrudescent parasites from new infections. However, these polymorphic genes are usually under selection. Therefore, neutral microsatellite markers are preferred as they are also easier to genotype. The current study investigated the genetic diversity and COI using the poly-α microsatellite marker to provide background information on circulating genotypes before its applied to TES in Kenya.

**Methodology:** Dried blood spot (DBS) samples were obtained from 93 participants from a TES in Busia County in 2016 and 92 participants from a malaria monitoring study conducted in Kilifi in 2020. Genotyping of the poly-α microsatellite was done by PCR, capillary electrophoresis and the fragment data analyzed using GeneMarker.

**Results:** About 96.7% and 87% of the samples from Busia and Kilifi, respectively, were successfully genotyped. The infections in Busia were mainly polyclonal (80%) with a significantly higher mean COI of 2.9 (p < 0.0001), while those in Kilifi were mostly monoclonal (52.5%) with a mean COI of 1.7. Despite on average a younger population and lower parasite density, both regions had similar expected heterozygosity (*He*) (Busia = 0.92; Kilifi = 0.90) while Busia recorded a slightly higher number of effective alleles (Ne) (Busia = 10.8; Kilifi = 9.3).

**Conclusion:** The poly-α microsatellite genotyping revealed high genetic diversity of malaria parasites in Busia and Kilifi. These findings define the genotypes (fragment sizes) observed in the two Kenyan populations, providing a proof of concept for the utility of poly-α in TES studies as a molecular correction tool and for the evaluation of the effectiveness of malaria interventions in Kenya.

## Introduction

Malaria continues to be a major public health burden worldwide. Despite the scaled up global efforts in combatting the ongoing disease transmission, there was a rise in malaria cases in 2023 to about 263 million cases and fortunately a decline in deaths to 597,000 when compared to 2022 that were 249 million cases and 608,000 deaths (WHO, 2024). Sub-Saharan Africa (SSA) is the most affected region with 95% of malaria cases and deaths being caused by *Plasmodium falciparum* (WHO, 2024). In Kenya, about 75% of the population is at risk of malaria infections with the Western Kenya and Coastal regions recording the highest community prevalence of about 18.9% and 4.5%, respectively (*Kenya National Bureau of Statistics. Kenya Malaria Indicator Survey, 2020*). Although the scale-up of control efforts has led to an improved coverage of interventions and reduced transmission, malaria is still problematic accounting for about 15% of the hospital visits in these regions (President’s Malaria Initiative, 2024).

Malarial control and eventual elimination are currently threatened by the presence of infections harboring numerous genetically distinct parasites, otherwise known as the complexity of infection (COI), and the presence of highly diverse *P. falciparum* parasites. The assessment of COI and genetic diversity of *P. falciparum* parasites is normally used to evaluate the impact of malaria control interventions as well as provide insights into parasite transmission dynamics (Apinjoh *et al*., 2019). The genetic diversity of the parasites emerges during the parasite life-cycle in the mosquito when genetic recombination occurs following the fusion of male and female gametes to form zygotes leading to the production of novel genotypes (Walliker *et al*., 1987; Li *et al*., 2019). Whereas the presence of multiple infections per individual (COI) happens either due to bites from a single mosquito carrying multiple parasite strains/clones or exposure to multiple infectious mosquito bites each introducing a different strain otherwise known as a superinfection (Nkhoma *et al*., 2020). For the parasite, the high genetic diversity improves its fitness to develop resistance to selective forces such as host immunity and drugs (Bose et al., 2016; Clay and Rudolf, 2019). A high COI within the host may enhance the development of multi-strain specific immunity resulting in a reduction in clinical symptoms and a controlling of parasitemia (Smith *et al*., 1999), while the high competition amongst the clones may increase gametocyte production and select for more virulent strains (Sondo *et al*., 2019), further complicating malaria control.

Currently, there are tools available for use to characterize the genetic diversity and COI of *P. falciparum* populations. These include molecular genotyping of highly polymorphic antigenic markers such as merozoite surface protein (*msp*) 1 and 2, and glutamate rich protein (*glurp*) using gel or capillary electrophoresis (Liljander *et al*., 2009). Genotyping these markers is recommended in anti-malarial drug therapeutic efficacy studies (TES) to differentiate recrudescent parasites from new infections (WHO, 2008). The analysis of *msp1* and *msp2* is tedious and complex by gel electrophoresis or GeneMarker, since they have 3 and 2 allelic family members, respectively and within each family there is extensive diversity based on the repeat regions under analysis. Thus, determining fragment sizes with accuracy requires extensive familiarity with the peak shapes and sizes, a clean and clear control panel and an understanding of the analytical settings. All this makes the fragment analysis pipeline complex. A potential alternative is the amplicon deep sequencing of apical membrane antigen (*ama*) 1 and other polymorphic markers with no repeat regions or shorter repeat regions, as well as whole genome sequencing techniques (Zhong, Lo, *et al*., 2018; Osborne *et al*., 2021; Wamae *et al*., 2022). Though the amplification methods and analytical pipelines require rigorous optimization of merozoite antigen markers that are subject to strong immune selection and are thus highly polymorphic, it provides a high-resolution quantitative method. The next-generation sequencing tools despite being highly sensitive in identifying minor genotypes especially in multiclonal infections, are still costly when the combination of reagents and run costs are considered in Africa (Zhong, Koepfli, *et al*., 2018). However, the use of these platforms is expanding across Africa that will enable a reduction in cost over time.

Recently, the use of microsatellites, short tandem repeats of 2-6 bp found throughout the *P. falciparum* genome, markers to study the parasite population structure has gained prominence (Touray *et al*., 2020; Ishengoma *et al*., 2024; Mwesigwa *et al*., 2024). These markers are highly polymorphic, easily amplifiable in the lab and are considered selection neutral since they are not under immune pressure (Schlötterer, 1998; Anderson *et al*., 1999; Zhong, Koepfli, *et al*., 2018). There are several microsatellites that are well characterized whose polymorphisms differ among parasite populations (Anderson *et al*., 2000). Studies have revealed that Poly-α, TAI, PfpK2 and TA42 are some of the most polymorphic microsatellites amongst parasite populations in the region (Gatei *et al*., 2015; Nderu *et al*., 2019; Touray *et al*., 2020; Mwesigwa *et al*., 2024). The World Health Organization (WHO) has also endorsed the use of these microsatellites to study parasite genetic diversity and has recommended that either Poly-α, Pfpk2 or TA1, should replace glutamate rich protein (*glurp)* that is considered less polymorphic and underestimates parasite genetic diversity, in anti-malarial drug efficacy studies (WHO, 2021).

Currently, there is insufficient knowledge on the diversity of poly-α microsatellite marker among the diverse parasite populations in Kenya to provide a background to the circulating genotypes in the population as this marker is introduced as a TES PCR correction marker. Therefore, the genetic diversity and COI of *P. falciparum* clinical infections was assessed from febrile cases attending primary healthcare facilities in two Kenyan populations, i.e. Busia in the Western Kenya region and Kilifi in the Coastal region (and to the East of Kenya), by genotyping the poly-α microsatellite marker.

## Materials and Methods

### Study design

This study used 93 Day 0 samples from Busia County (a high malaria transmission region) collected from a TES that was conducted between August and September 2016 (Nderu *et al*., 2019) and 92 samples collected from a malaria monitoring study collecting dried blood spots (DBS) samples from microscopy and rapid diagnostic test (RDT) confirmed malaria cases that was conducted between December 2019 and February 2020 in Kilifi County (a moderate to high malaria transmission region). In the TES study, children between the age of 6 months to 14 years with confirmed malaria infection (by microscopy) were randomized to receive dihydroartemisinin-piperaquine (DP) or artemether-lumefantrine (AL). The parasite density ranged from 1,000 to 100,000 in Busia and 160 to 720,000 in Kilifi. Written informed consent was obtained from the guardians of the study participants prior to sample collection. Ethical approval was obtained from the Kenya Medical Research Institute’s (KEMRI) Scientific Ethics and Review Unit (SERU) under protocol number 3403 for the TES study and 2617 for the malaria monitoring study.

### DNA extraction

Samples from the two locations were accessed from the biobank in May 2023. Parasite genomic DNA was extracted using QIAcube HT according to the manufacturer’s instructions for the Kilifi samples and by the Chelex extraction method for the Busia samples. DNA extraction by Chelex- saponin method was performed as follows: Each DBS was punched to two 2.5-mm discs with a sterile (absolute ethanol [96%] and a flame) puncher at the center and periphery and transferred to a 1.5mL Eppendorf tubes with sterile tweezers. The samples were lysed overnight using 1mL of 0.5% (w/v) saponinin/1X phosphate-buffered saline (PBS). Following saponin aspiration, the discs were incubated in 1mL 1X PBS at 4°C for 30 min; thereafter, 150mL of a solution of 6% (w/v) Chelex in DNase/RNase-free water was used to incubate the samples for 30 min at 97°C. At regular (10min) intervals, the samples were vortexed and centrifuged to maximize the elution of DNA. The plates were then centrifuged at 4,000 × g for 5 min, and 120μl of the DNA-containing solution was stored at −20°C for further analyses.

### Genotyping of samples

A 15μl PCR assay was conducted to amplify the poly-A marker. Fluorescently labelled primers were used (forward, 5‘- AAAATATAGACGAACAGA-3‘; reverse, 5‘- ATCAGATAATTGTTGGTA -3’ FAM). The expected size range was 100 – 201 based on (Anderson *et al*., 2000). The reaction mix included 7.5μl of (Promega, Catalogue number: M7505), 0.6μl each of forward and reverse primers, 7.5μl nuclease-free water and 2μl of template DNA. Cycling conditions were as follows: initial denaturation at 94°C for 2 min, followed by 25 cycles of denaturation at 94°C for 20 sec, annealing at 45°C for 20 sec, extension at 65°C for 30 sec, and a final extension at 65°C for 2 min. The PCR products were visualized on 1.5% (w/v) agarose gel electrophoresis (Sigma-Aldrich, USA) in 0.5X TBE buffer. After staining with Redsafe® (iNtRON Biotechnology, Korea), the fragments were visualized under UV light using a Universal Hood II (Bio-Rad Laboratories, Inc., USA). Fragment sizes were estimated by comparison to a 100 base-pair DNA ladder (Promega, USA). About 1.5μl of the successfully amplified samples were mixed with 9 μl of Hi-Di formamide, and 0.5μl of size standard (ROX-350, Applied Biosystems). The solutions were transferred to 96-well Optical reaction plates and sent to the International Livestock Research Institute (ILRI) in Nairobi, Kenya, for capillary electrophoresis on the 3730xl DNA sequencer (Applied Biosystems).

### Fragment data analysis

Analysis of the chromatograms was carried out using GeneMarker™ software version 3.0.1 (SoftGenetics). A panel was first created based on poly-α microsatellite size range that was adopted from (Anderson *et al*., 2000). A relative fluorescence unit (RFU) threshold of 1000, based on the negative control, was used to distinguish true peaks from background noise and non-specific peaks.

Major peaks above the RFU threshold and minor alleles with at least 30% of the major allele’s height were also scored. All chromatograms were manually reviewed to confirm the peaks. Fragment sizes were binned into 3 nucleotide windows. Since *P. falciparum* parasites are usually haploid, each genotype detected represented a distinct clone of the parasite. Consequently, the number of clones per individual infection in each population was defined as the complexity of infection (COI). The mean COI per population was calculated by dividing the total number of genotypes detected by the number of samples in that population. Mann–Whitney U test was used to compare continuous variables, i.e., COI, parasitemia and age, between the two populations, a p-value less than 0.05 was considered statistically significant. Isolates were categorized as monoclonal if they had a single genotype and polyclonal if they had at least two genotypes. For polyclonal samples, only the major genotype was used in the analyses of the expected heterozygosity (*He*). The *He* was described as the probability that two genotypes selected at random from a population will contain different alleles and ranges from 0 (no allele diversity) to 1 (all alleles are different). *He* was used to estimate poly-α allelic diversity at each location based on the formula below.

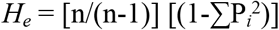

where n is the sample size and P*i* is the frequency of *i*th allele in the population (Nei, 1978).

The metadata for the samples was accessed on 30^th^ May 2023 for the Kilifi samples and 30^th^ October 2024 for the Busia samples. Data analysis was performed using R version 4.4.2 (R Core Team, 2023) and graphical outputs generated using ggplot2 (version 3.5.1) package (Kassambara, 2020).

## Results

### Characteristics of study participants

Among the 93 samples screened in Busia in 2016, 86 had patient data while in Kilifi in 2020, 71 samples had patient data out of 92 samples. The participants from Busia were primarily female (53%) while those from Kilifi were mainly male (61%). Participants from Kilifi (median age = 5.3; IQR 3.0 – 8.3) were older than those from Busia (median = 3.8; IQR= 2.0 – 6.2). This age difference was statistically significant, *P* = 0.0174. The Kilifi participants also harbored significantly higher infections with a significantly high median parasitemia of150,000 (IQR: 7120, 265,000.0) compared to Busia median = 22,963 (IQR: 5,680.0, 37,480.0) based on microscopy (*P*=0.001).

### Complexity of *P. falciparum* infections (COI)

In malaria endemic regions, individuals often carry multiple genetically distinct parasite clones otherwise known as complexity of infection (COI). Individuals with one poly-α genotype were categorized as monoclonal, while those with more than one genotype were referred to as polyclonal. The majority of individuals in Busia had a high mean COI of 2.9 compared to Kilifi, COI = 1.7 (**Figure 1A**) this difference was statistically significant (p < 0.0001). Similarly, 80% of the infections in Busia were polyclonal, while in Kilifi these infections were lower at 47.5% (**Figure 1B**), with the highest number of clones per infection being 7 and 4, respectively.

**Figure 1:**
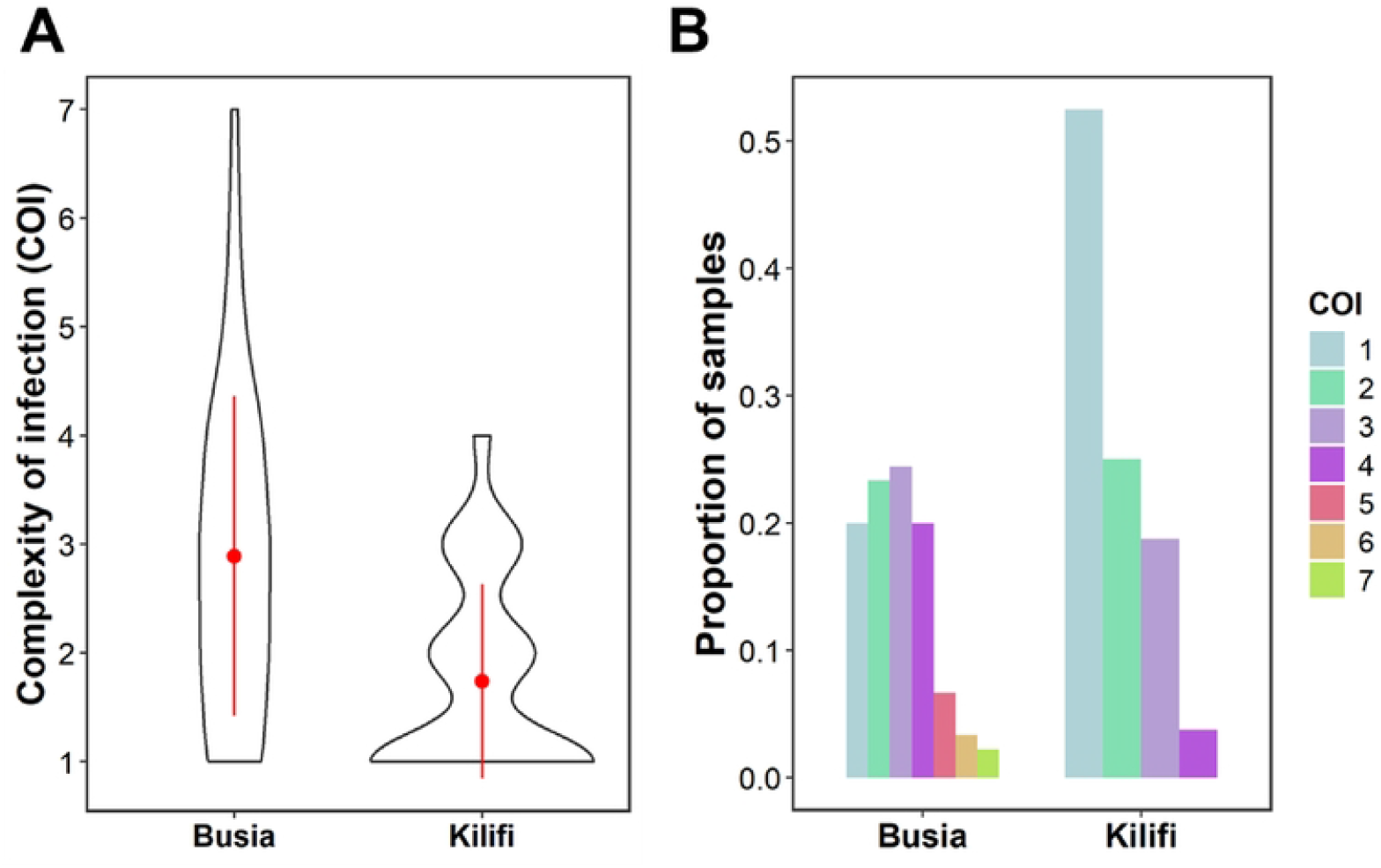
Complexity of infection (COI) across the two locations. **A)** Distribution of COI in Busia and Kilifi. The mean COi is depicted by the red dot and the red vertical line shows one standard deviation away from the mean. B) Proportion ofsrunples with genetically distinct genotypes detected in the uvo locations. The different colors represent the different number ofclones as sho,vn in the key.

### Genetic diversity of *P. falciparum* infections based on poly-α

A total of 31 poly-α genotypes were reported from the two populations that ranged in size from 103bp to 210bp. Nineteen of these genotypes were prevalent in both locations, while five genotypes (102bp, 105bp, 111bp, 114bp and 195bp) were exclusively reported in Kilifi and seven genotypes (108bp, 126bp, 138bp, 177bp, 180bp, 186bp and 189bp) were exclusively reported in Busia. About 26 distinct genotypes (Na) were revealed from the 90 isolates that were successfully genotyped from Busia. These genotypes ranged in size from 103 to 210bp. The number of effective alleles (Ne) was 16.1 (**Table 2**). The most common genotypes from Busia were 150bp, 153bp and 156bp that recorded frequencies of 11.2%, 9.6% and 10.8%, respectively (**Figure 2**). In Kilifi, 24 distinct genotypes, ranging in size from 102 to 210bp, were revealed from the 80 successfully genotyped samples (**Table 2**). Out of these, the three most common genotypes were 150bp, 153bp and 210bp whose frequencies were 14.4%, 18.7% and 11.5%, respectively (**Figure 2**). The 156bp fragment was the 4^th^ common genotype in Kilifi. The 210bp fragment is unique with a very high frequency in the Kilifi population. In both populations, Ne (Busia = 10.8; Kilifi = 9.3), that represents the number of unique genotypes that would need to be equally frequent to achieve the same level of genetic diversity, was lower than Na indicating that majority of the genotypes occurred at low frequencies. The *He*, was similar in Busia (0.92) and Kilifi (0.9) (**Table 2**).

**Figure 2:**
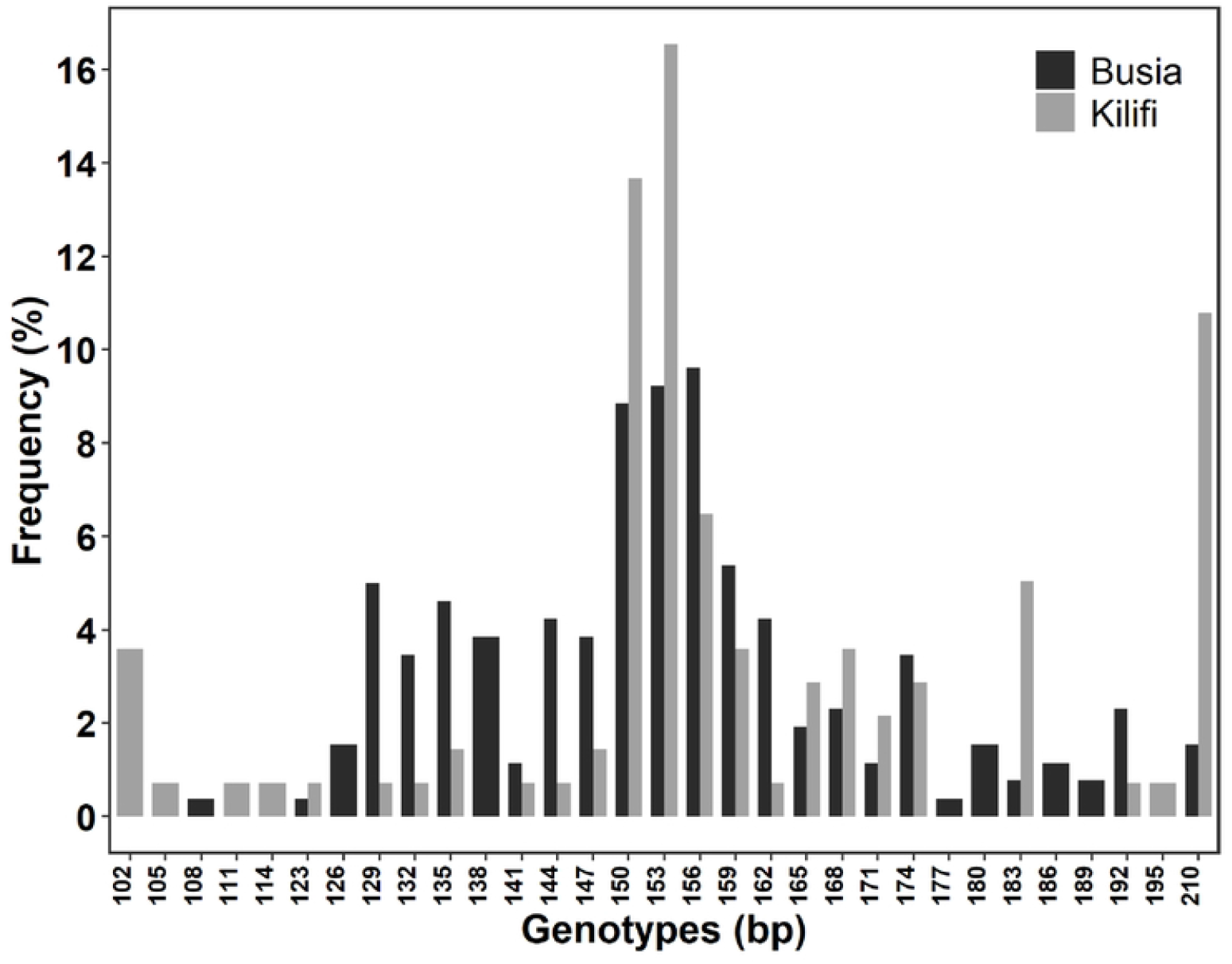
Frequencies of poly α alleles in Busia (black) and Kilifi (gray). The horizontal axis shows the individuals alleles sizes in base pairs (bp), vhile the vertical axis shows their respective frequences in percentage.

## Discussion

The poly-α microsatellite genotyping revealed infections with high polyclonality and mean COI in Busia compared to Kilifi in low parasitemia infections, which could be attributed to the higher transmission intensity in Busia and the younger population sampled. This is consistent with other studies conducted in these regions revealing polyclonality levels between 75% - 80% in Western Kenya (Nderu *et al*., 2019; Touray *et al*., 2020) and 40% - 50% in Coastal Kenya (Nderu *et al*., 2019; Kimenyi *et al*., 2022) using microsatellites and other polymorphic markers. A high COI of 2.9 in Busia and 1.7 in Kilifi is also comparable to other studies in these regions that reported a mean COI of 3.4 in Mbita, Western Kenya (Touray *et al*., 2020) and 1.6 in Msambweni, Coastal Kenya (Nderu *et al*., 2019). The average parasite density in Busia was almost six-fold lower than Kilifi alluding to a high anti-disease immunity in the older population in Kilifi who would have experienced multiple malaria exposures. Thus, to experience malaria symptoms a higher parasite burden is required to overcome the developed malaria immunity over time.

The high polyclonality of infections is further corroborated by the higher *He* in Busia complemented by the higher total number of genotypes, that is expected from a high transmission region. Other studies that have characterized the poly-α microsatellite marker have also revealed almost similar findings i.e. Mbita (*He* = 0.95) (Touray *et al*., 2020), Busia (0.92), Nyando (0.88) (Nderu *et al*., 2019), Asembo (*He* = 0.91) and Gem (*He* = 0.89) (Gatei *et al*., 2015) in Western Kenya and Msambweni (*He* = 0.85) in Coastal Kenya (Nderu *et al*., 2019). The high genetic diversity in Western Kenya may also be attributed to importation of diverse parasite strains from neighboring malaria endemic areas (Mulenge *et al*., 2016). Thus, elucidating the impact of malaria control interventions using information from parasite genetic diversity studies should account for external factors impacting parasite population genetics. The most common poly-α genotypes included the 150 bp and 153 bp genotypes in Busia and Kilifi, respectively. In 2019, a Kenyan study reported that the 153 bp genotype was the most prevalent in Busia and Nyando while the 171 bp genotype was the most prevalent in Msambweni (Nderu *et al*., 2019). Likewise, a study in Uganda has recently reported that the 153 bp genotype was the most prevalent among asymptomatic, uncomplicated and complicated malaria infections (Mwesigwa *et al*., 2024). The 210bp fragment is unique with a high frequency in the Kilifi population, based on the assessment of the fragment sizes in GeneMarker it appears to be a true fragemnt that requires further validation in future studies. The analysis of poly-α was simple since it does not have any allelic family members and a shorter repeat region while still having the capability to define COI and several alleles within and between populations.

This study based its findings on one microsatellite marker which may underestimate the parasite genetic diversity as it may fail to capture the entire genome-wide genetic diversity. Thus, the use of several microsatellite markers is recommended. However, it will require an analytical algorithm to determine with high accuracy the diversity in the population. Furthermore, it has been shown that genotypes/fragments may converge to similar sizes while not having identical sequence composition (Takala *et al*., 2006), leading to underestimation of diversity. Thus, sequencing provides a higher resolution approach to decipher genetic diversity. The amplicon next-generation sequencing approach provides both qualitative and quantitative data to assess parasite population genetic diversity at high resolution to better determine COI (Lerch *et al*., 2017). Though there are differences in the timing of samples collections and geographical locations, any genetic differences could be due to either time or geography. For example, the 210 bp fragment observed primarily in Kilifi compared to Busia would require additional sampling from the same time points from other regions in the country to resolve the differences observed. Nevertheless, the poly-α data provided a representation for circulating fragments sizes in Kenya and as a single marker indicated much lower malaria transmission in Kilifi compared to Busia based on the lower COI and more than half the infections being monoclonal. It is thus capable of distinguishing new from recrudescent infections in TES in addition to other polymorphic markers.

## Acknowledgements

We thank the study participants as well as their parents/guardians for giving consent. We also extend our gratitude to Naomi Lucchi from the President’s Malaria Initiative (PMI) in Rwanda for her training on the poly-α PCR assay. This manuscript was written with the permission of Director KEMRI CGMRC.

## Author Contributions

LIO conceptualized the study, NA, conducted the assays, LN, LIO and NA optimized the data analysis pipeline, CM, NA, LN and KMK conducted the analysis and NA, KMK and LIO drafted the manuscript. All authors contributed and reviewed the final manuscript.

## Data Availability

The datasets supporting the conclusion of this article are available in the Harvard Dataverse repository: https://doi.org/10.7910/DVN/DAHYSD.

